# Comparisons of ribonuclease HI homologs and mutants uncover a multi-state model for substrate recognition

**DOI:** 10.1101/2021.11.04.467286

**Authors:** James A. Martin, Arthur G. Palmer

## Abstract

Ribonuclease HI (RNHI) non-specifically cleaves the RNA strand in RNA:DNA hybrid duplexes in a myriad of biological processes, including retroviral reverse transcription. Several RNHI homologs contain an extended domain, termed the handle region, that is critical to substrate binding. Nuclear magnetic resonance (NMR) spectroscopy and molecular dynamics (MD) simulations have suggested a kinetic model in which the handle region can exist in open (substrate-binding competent) or closed (substrate-binding incompetent) states in homologs containing arginine or lysine at position 88 (using sequence numbering of *E. coli* RNHI), while the handle region populates a state intermediate between the open and closed conformers in homologs with asparagine at residue 88 [Stafford, K. A., et al., *PLoS Comput. Biol.* **2013**, 9, 1-10]. NMR parameters characterizing handle region dynamics are highly correlated with enzymatic activity for RNHI homologs with two-state (open/closed) handle regions [Martin, J. A., et al., *Biochemistry* **2020**, 59, 3201-3205]. The work presented herein shows that homologs with one-state (intermediate) handle regions display distinct structural features compared with their two-state counterparts. Comparisons of RNHI homologs and site-directed mutants with arginine at position 88 support a kinetic model for handle region dynamics that includes 12 unique transitions between eight conformations. Overall, these findings present an example of the structure-function relationships of enzymes and spotlight the use of NMR spectroscopy and MD simulations in uncovering fine details of conformational preferences.

## INTRODUCTION

Ribonuclease HI (RNHI) is an omnipresent, non-sequence-specific, endonuclease that cleaves the RNA strand in RNA:DNA hybrid duplexes. The enzyme has roles in replication,^1^ genome maintenance,^2^ and is the C-terminal domain of retroviral reverse transcriptase (RT) proteins.^3–4^ Murine Leukemia Virus (MLV) and Human Immunodeficiency Virus (HIV) are two such retroviruses and their RNHI domains are necessary for viral replication.^5–6^ A number of RNHI homologs contain a “handle region”, an extended loop (residues 81-101, numbering based on *E. coli* RNHI is used herein) with a large cluster of positive residues, that is critical for substrate recognition (Figure 1A-B).^8^ Solution-state nuclear magnetic resonance (NMR) spectroscopy^9^ combined with Molecular Dynamics (MD) simulations^10^ have suggested a two-state kinetic model in which the handle region of the mesophile *E. coli* RNHI (EcRNHI) populates “open” (substrate-binding competent) and “closed” (substrate-binding incompetent) states, while the thermophile *Thermus thermophilus* RNHI (TtRNHI) predominantly populates the closed state at 300 K (Figure 1C).

**Figure 1.**
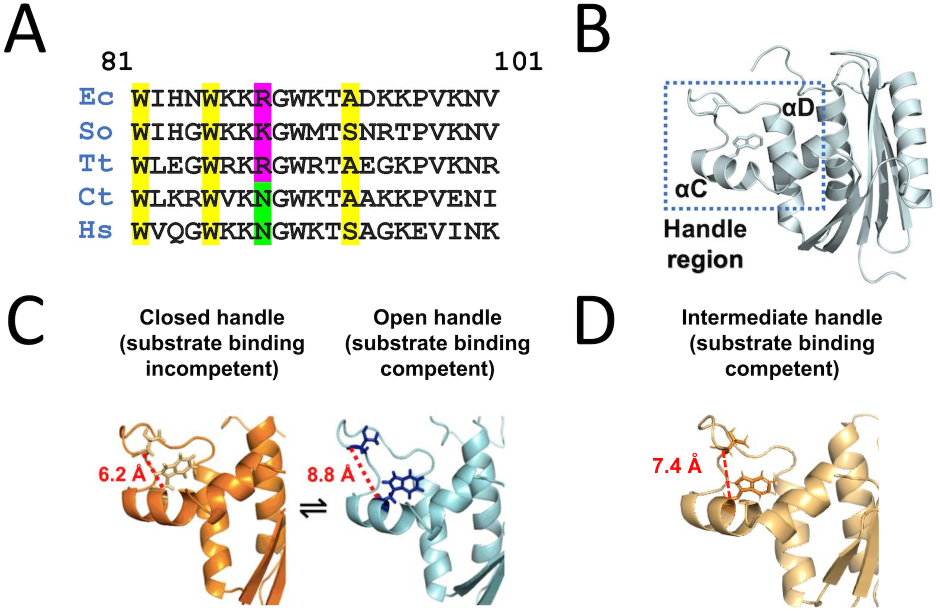
Ribonuclease HI Handle Region. **(A)** Multiple sequence alignment of RNHI handle regions from *E. coli* (Ec), *S. oneidensis* (So), *T. thermophilus* (Tt), *C. tepidum* (Ct), and *H. sapien* (Hs). Yellow highlights: DNA-binding residues.^11^ Sequence numbering is according to EcRNHI. **(B)** Tertiary structure of EcRNHI with a focus on the handle region. Trp 85 (W85) and Thr 92 (T92) are shown in sticks. **(C)** Two-state kinetic model for handle region dynamics.^10^ Open and closed states are characterized by the distance from the Cα of W85 that forms part of a DNA-binding channel^11^ to the Cα of T92 at the tip of the dynamic handle loop; open distances are taken to be ≥ 7.8 Å and closed distances < 7.8 Å. Homologs with Lys or Arg at residue 88 (pink) follow this model; those with Asn (green) display a handle region conformation intermediate between the open and closed conformers (**D**).

This model has been extended by correlating NMR parameters describing the handle region conformation, including residue 101 χ1 trans percentage and Trp 85 Nε1-Hε1 and Thr 92 N-H residual dipolar couplings (RDCs) with enzymatic activity for each homolog, in what is termed the Weighted Conformer Model.^12^ However, this model was developed only for homologs with handle regions exhibiting two-state dynamics and able to populate open and closed conformations. Homologs that populate a single conformation intermediate between the open and closed conformations have been identified through MD simulations (Figure 1D).^10^ Two-state (open/closed) or one-state (intermediate) behavior of the handle region arises from the identity of residue 88: long side-chain, positive amino acids (lysine and arginine) are associated with the former, while the shorter, neutral amino acid asparagine is associated with the latter. EcRNHI, TtRNHI, and the psychrophile *Shewanella oneidensis* RNHI (SoRNHI) have two-state handle regions. The moderate thermophile *Chlorobium tepidum* RNHI (CtRNHI) and the *Homo sapiens* RNHI domain (HsRNHI) have one-state handle regions. HsRNHI is the only RNHI homolog containing a handle region that has been co-crystallized with substrate;^11^ apo-EcRNHI residues K86, K87 and R88 form crystal contacts with acidic active sites of neighboring molecules.^13–14^

The present work investigates whether homologs with one-state handle regions display dynamic and enzymatic properties that can be characterized similarly as for two-state counterparts. NMR experiments characterizing handle region dynamics were performed as described previously.^12^ Reciprocal mutations were made in EcRNHI (R88N) and CtRNHI (N88R) to determine the effect of the residue type at position 88. Further comparisons between RNHI homologs were made using a combination of Circular Dichroism (CD) spectroscopy, NMR spectroscopy, MD simulations, and x-ray crystallographic structures. These data establish N88 as an important contributor in the thermostability of the hydrophobic core of RNHI, as well as amend the kinetic model for handle region dynamics to include 12 unique transitions between eight total conformations.

CtRNHI is a moderate thermophile with 57% sequence identity to TtRNHI and 53% sequence identity to EcRNHI. Its overall stability and melting temperature are similar to EcRNHI, but its change in heat capacity upon unfolding and stability at optimal growth temperature are more similar to TtRNHI. Finally, its optimal growth temperature is intermediate EcRNHI and TtRNHI.^15^ The x-ray crystal structure of CtRNHI has been solved at a resolution of 1.60 Å (PDB ID: 3H08) and shows two monomers in the unit cell, both with missing residues in the handle region (Figure S1).^15^ Superposition of CtRNHI and TtRNHI shows higher structural similarity of the handle regions (0.754 Å Cα-RMSD) compared to EcRNHI (0.975 Å Cα-RMSD). CtRNHI has a characteristic handle distance of 6.7 Å (see Fig.1 caption), which is more similar to TtRNHI (6.5 Å) than EcRNHI (7.8 Å). However, structures of CtRNHI observed in MD simulations have handle distances (~7.4 Å) that are intermediate and more like distances in EcRNHI.^10^ The disparity between the two measurements provides ambiguity about the handle region dynamics of CtRNHI.

## RESULTS

Near complete backbone and sidechain NMR resonance assignments were made for CtRNHI (all NMR experiments herein used Cys-free mutant proteins) and handle region dynamics were characterized by NMR spectroscopy as described previously.^12^ The Weighted Conformer Score calculated for CtRNHI from these NMR parameters (0.31) is intermediate between EcRNHI (0.21) and TtRNHI (0.63) (Table S1). If CtRNHI has handle region dynamics similar to homologs with two-state handle regions, then the weighted conformer score predicts a Michaelis constant (K_M_) that is approximately 2.4 (± 0.4) fold greater than EcRNHI (Figure 2A). In contrast, enzymatic assays for CtRNHI and EcRNHI yield a relative Michaelis constant of 0.77 ± 0.29 (Figure 2B). These data suggest that the contributions of handle region dynamics to substrate recognition by CtRNHI differ in some regards to two-state handle region RNHI homologs.

**Figure 2.**
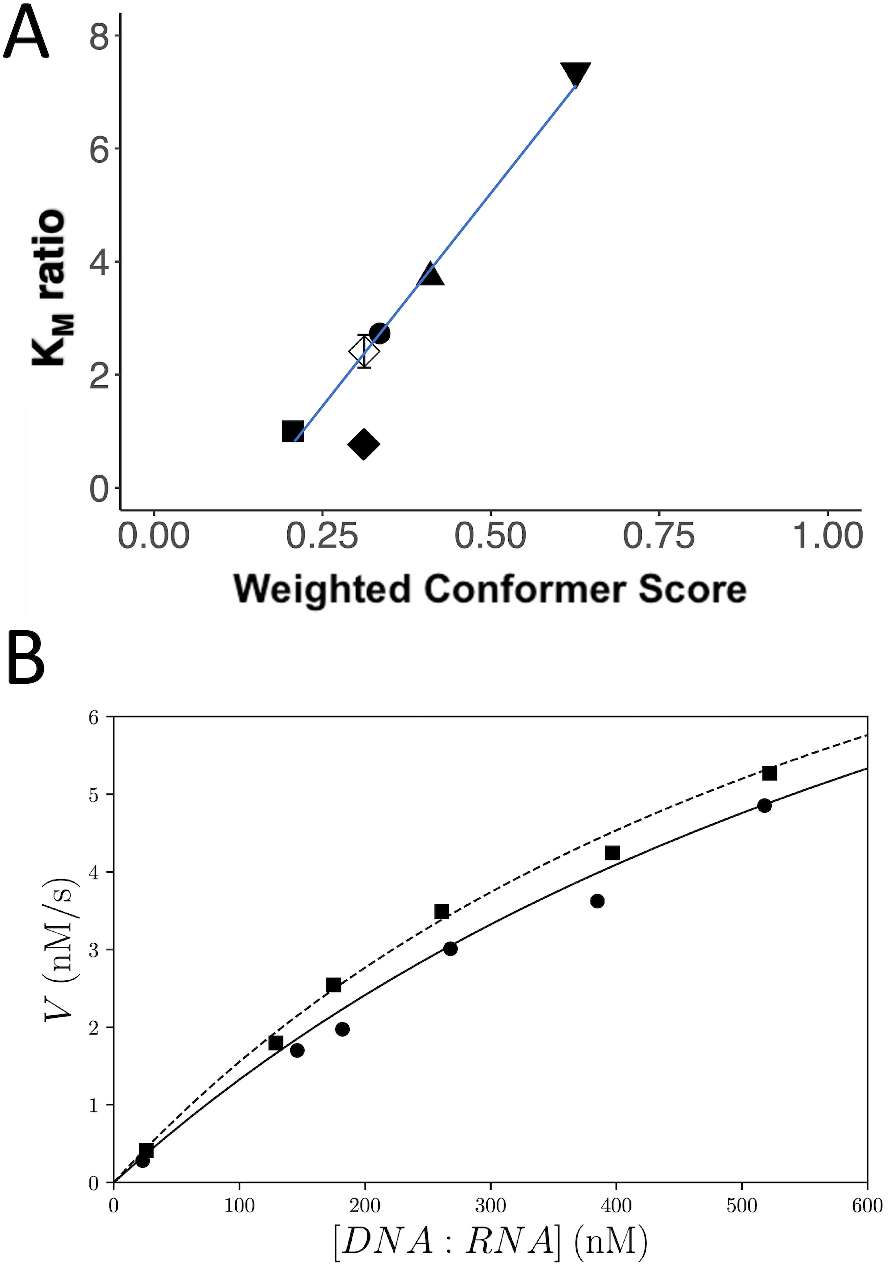
CtRNHI Enzyme kinetics. **(A)** *Open Diamond*: Michaelis constant (K_M_) prediction for CtRNHI using the Weighted Conformer Model.^12^ Experimental K_M_ ratios: K_M_ for each RNHI homolog normalized by the corresponding K_M_ for EcRNHI. ***Square***: EcRNHI, ***Circle***: SoRNHI, ***Triangle***: V98A EcRNHI, ***Inverted Triangle***: TtRNHI, ***Closed Diamond***: CtRNHI **(B)** Initial reaction velocities are plotted versus substrate concentrations for EcRNHI (circle, solid line) and CtRNHI (square, dashed line). Lines are calculated from the Michaelis-Menton equation using values of K_M_ and V_max_ obtained from global fits of progress curves. Fitted values of K_M_ and V_max_ are 920 ± 240 nM and 13.5 ± 3.2 nM/s for EcRNHI and 710 ± 190 nM and 12.6 ± 2.9 nM/s for CtRNHI, respectively.

Reciprocal mutations were made in CtRNHI (N88R) and EcRNHI (R88N) to better understand the features of one-state handle regions. Backbone and sidechain NMR resonance assignments were obtained for each mutant and compared to its respective wildtype counterpart. Overlaid ^15^N and ^13^C heteronuclear single quantum coherence (HSQC) NMR spectra for CtRNHI show that the structural changes caused by the mutation are localized to the handle region (Figure S2). Specifically, resonances for K91 (the interaction partner of N88 in wild-type CtRNHI), V98 and I101 are the most perturbed, consistent with previous research suggesting that these residues have concerted movements.^12^ K91 is also most perturbed in R88N EcRNHI, however, the overall differences are small and broadly distributed (Figure S3, consistent with studies showing that dynamics of the handle region are correlated with structural elements throughout EcRNHI.^16–17^ The NMR chemical shift perturbations are complemented by nearly identical CD spectra for the respective homolog/mutant pairs, suggesting the secondary structure of the enzyme remains unchanged upon mutation (Figure 3A-B). Thermal denaturation is more reversible for wild-type and N88R mutant CtRNHI than for wild-type and R88N EcRNHI; the thermal melts also show an 11% decrease in reversibility for N88R CtRNHI (Figure 3C) and a 9% increase in reversibility for R88N EcRNHI (Figure 3D).

**Figure 3.**
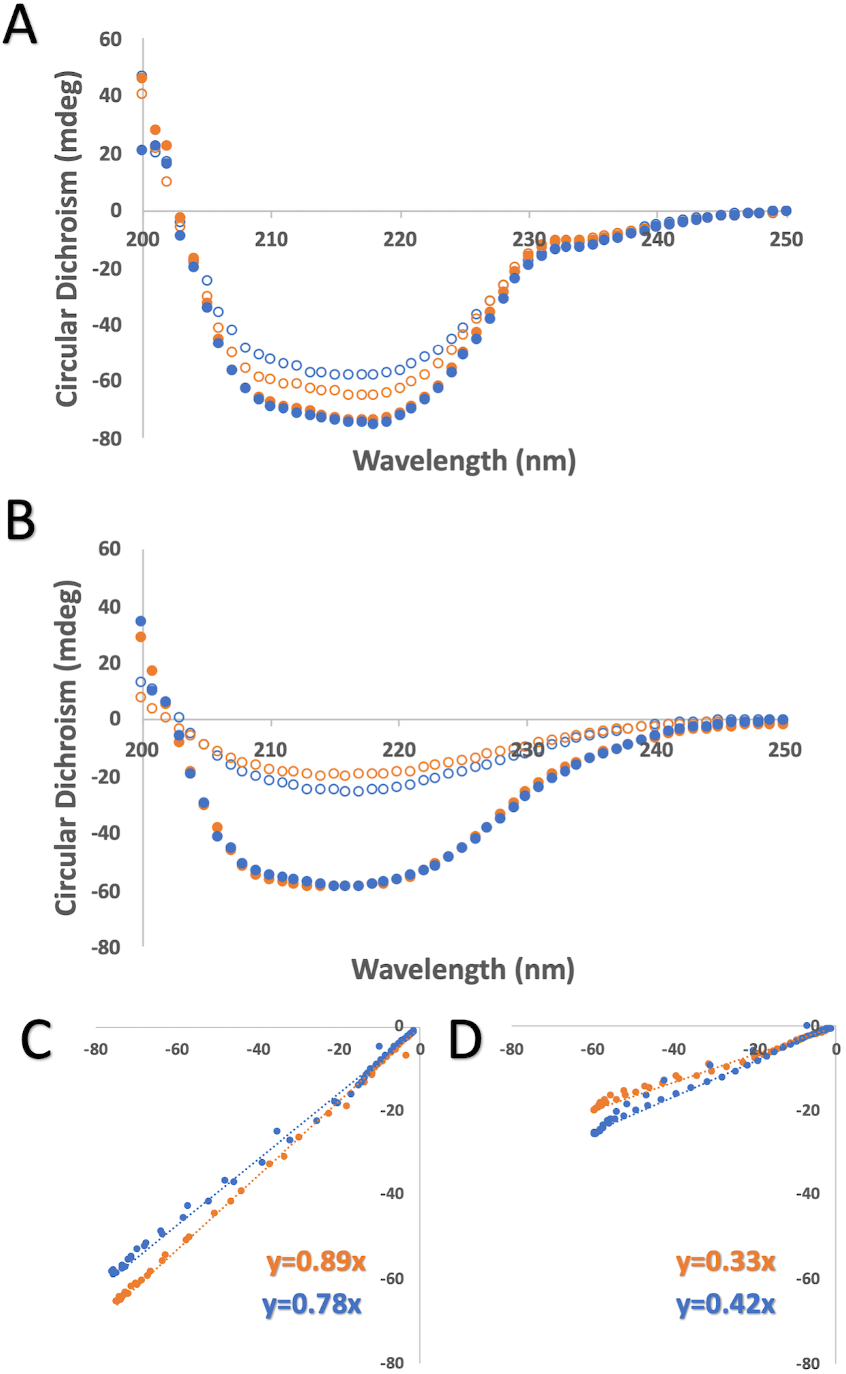
Far-UV CD Spectra of wild-type and mutant RNHI enzymes. CD spectra recorded at 25°C for **(A)** *Orange:* CtRNHI, *Blue:* N88R CtRNHI and **(B)** *Orange:* EcRNHI, *Blue:* R88N EcRNHI proteins before (closed circles) and after (open circles) denaturation at 85 °C. CD signal from native (x-axis) vs renatured (y-axis) enzymes are respectively plotted for **(C)** CtRNHI and **(D)** EcRNHI. Slopes represent the recovery of CD signal (reversibility) over the spectra range.

Experimental NH RDCs, an NMR parameter used for structure validation and refinement,^18^ agree poorly with values fitted to the crystal structure of CtRNHI (Figure S4A), with a Q factor of 0.50 and R = 0.81; agreement is slightly worse when only handle region residues are considered, with R = 0.76 (Figure S4B). Thus, the structure of the backbone and handle region in the crystal structure of CtRNHI may differ from the structure(s) in solution. MD simulations for CtRNHI were used to obtain a structural model more representative of the experimental RDCs. Briefly, experimental RDCs were fit to individual structural frames of a 100 ns trajectory at 300 K to identify the structure (BF CtRNHI) with the lowest Q factor.^18^ Agreement improved in comparison to the crystal structure for backbone NH RDCs (Figure S4C), with Q = 0.37 and R = 0.85, and for the handle region (Figure S4D) with R = 0.89. Additionally, the *in silico* structure BF CtRNHI has a better Molprobity score (metric evaluating the quality of a structure)^19^ than either of the monomers in the crystal structure (Table S2).

The handle distance for BF CtRNHI is 7.2 Å, intermediate between EcRNHI (7.8 Å) and TtRNHI (6.5 Å). Inspection of the handle region of BF CtRNHI (Figure 4A) shows that the side chain of N88 makes two hydrogen bonds with the backbone of K91: N88 Oδ1:H K91 and N88 Hδ2:O K91 (as noted previously)^10^ and none with T92. HsRNHI is the other homolog that naturally contains asparagine at residue 88. Superposition of BF CtRNHI and HsRNHI (2QK9) show similar architecture, even though the HsRNHI structure was determined in complex with substrate (Figure 4B). The handle regions of HsRNHI and BF CtRNHI have a Cα-RMSD of 0.65 Å. The main perturbations occur at the end of the handle loop (after V98). When the end of the handle loop is excluded from alignment, the Cα-RMSD reduces to 0.47 Å. One of the hydrogen bonds (N88 Hδ2:O K91) is longer in HsRNHI (2.2 Å vs 2.0 Å), and the distance between N88 Nδ2-Hδ2 and T92 O is also larger (5.1 Å vs 4.9 Å). These slight differences combined with the steric constraint due to the presence of substrate may account for why HsRNHI is more open (7.8 Å) than BF CtRNHI (7.2 Å).

**Figure 4.**
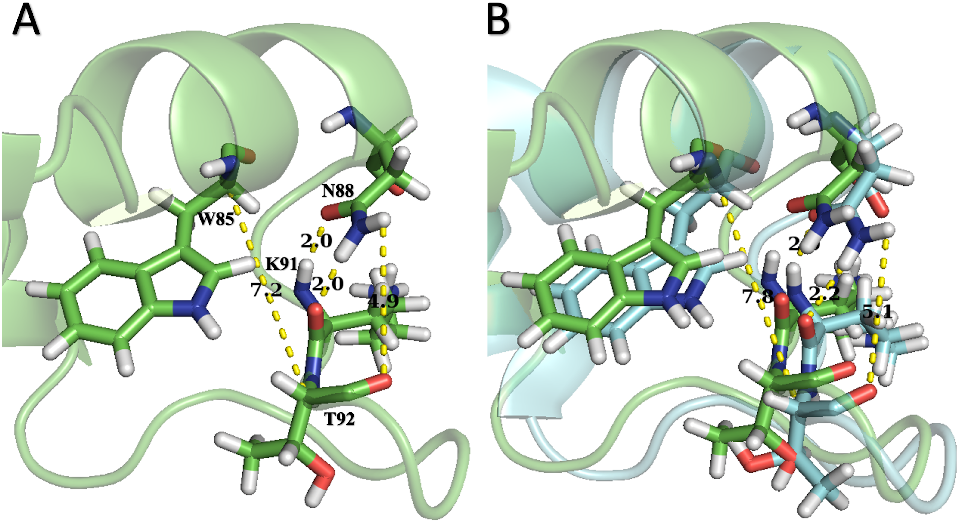
N88 Interactions of HsRNHI and CtRNHI. **(A)** Closeup of the handle region of BF CtRNHI with a focus on W85, N88, K91 and T92. **(B)** Overlay of HsRNHI(cyan) and BF CtRNHI (green) handle regions. The overall architecture of the interactions of N88 with K91 and T92 remains the same. Sequence numbering is according to EcRNHI.

Structural analysis of a sextuple, thermostable mutant of SoRNHI (6X SoRNHI)^20^ offers further insight into the interactions of N88 with other handle region residues. 6X SoRNHI has six mutations: N27K, D37G, M74V, K88N, R95G, and D134H that increase its thermostability by 28.8 K and decrease its relative activity to 43% of the wildtype enzyme. The effects of the mutations on stability are approximately additive.^20^ Furthermore, the overall structure of 6X SoRNHI (2ZQB) is unaltered outside of the mutation sites. The four monomers in the crystallographic unit cell (Figure 5A-D) are structurally similar to each another (Figure 5E) and have an all Cα-RMSD to wild-type SoRNHI in the range of 0.653-1.062 Å (Table S3). The handle region for each monomer is also structurally similar to the wildtype structure, although the position of the handle loop relative to the protein core differs between monomers (vide infra).^20^

**Figure 5.**
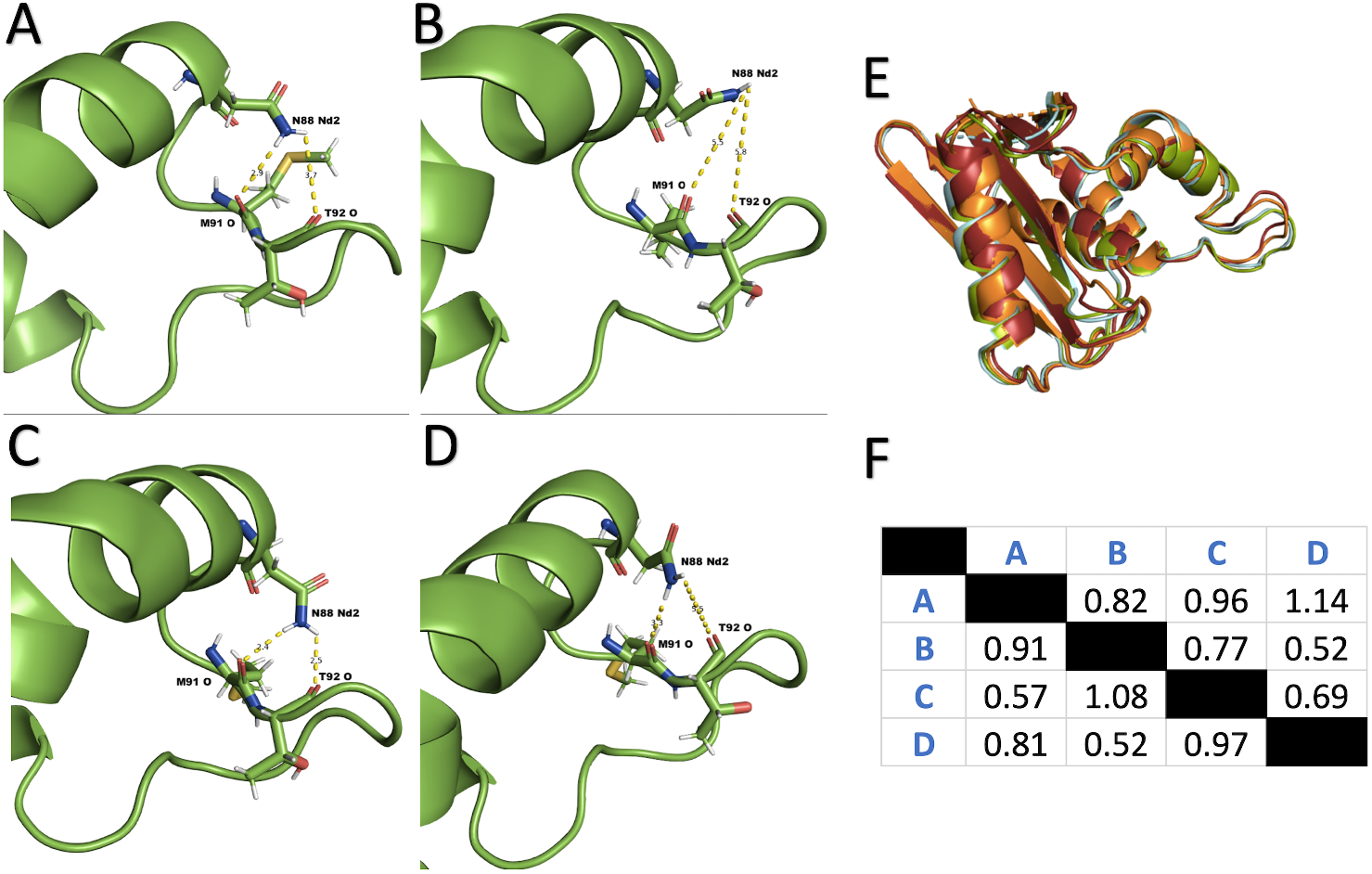
N88 Interactions and structural comparisons of 6X SoRNHI monomers. Handle region of each monomer, A-D (labeled accordingly), of 6X SoRNHI (2ZQB). (E) Overlay of all 4 monomers of 6X SoRNHI: Monomer A (orange), B (cyan), C (red), and D (green). (F) Matrix of Cα-RMSDs (Å) for the overall structure (above diagonal) and for the handle region (below diagonal).

These observations suggest the mutations act independently of one another. This conclusion is consistent with a study of a quintuple thermostable EcRNHI mutant (5X EcRNHI: G23A, H62P, V74L, K95G, D134H) that showed the structures at the mutation sites are almost identical to the respective structures for the single mutant variants.^21^ Thus, with reasonable certainty, the interactions of N88 in 6X SoRNHI are primarily the effect of the K88N mutation.

The interactions of N88 in the sextuplet mutant mirrors that of the other homologs and variants containing N88. The N88 sidechain interacts with the backbones of M91 and T92 in monomers A and C; according to the handle distance metrics (Table S3), their handle regions would be considered closed (6.8 and 6.7 Å respectively). The N88 side chain points away from the backbones of M91 and T92 in monomer B; its handle region metrics would classify monomer B as the most open structure (7.6 Å). The N88 side chain interacts with M91 O but not with T92 O in monomer D. This conformation is nearly identical to the other three monomers, with the main difference being the hydrogen bond with M91 O is longer and N88 Oδ1 does not form a hydrogen bond with M91 NH. Nonetheless, Monomer D appears to be in a transition state from closed to open; its handle metrics are characteristic of a closed structure, but its handle distance (7.4 Å) is closer to that of Monomer B (7.6 Å) than A and C (6.8 and 6.7 Å respectively).

A matrix of Cα-RMSDs for the four monomers show that monomers B and D are most alike overall (Figure 5F, upper triangle) and in their handle regions (Figure 5F, lower triangle). The handle conformations of monomers A and C are much more similar than to either monomers B and D (Fig. 5F, lower triangle). Due to the high overall structural similarity of the four monomers, each monomer may represent a step in the transition of the handle region from closed to open. The process may start with the most closed monomer (C: 6.7 Å) stabilized by hydrogen bonds between N88 Nδ2-Hδ2 and M91 O (2.4 Å) and T92 O (2.5 Å). Next, these hydrogen bonds lengthen (2.9 Å and 3.7 Å respectively) as the handle region becomes slightly less closed (Monomer A: 6.8 Å). The hydrogen bond between N88 Nδ2-Hδ2 and T92 O breaks (5.5 Å), transitioning the structure from closed to open (Monomer D: 7.4 Å). Lastly, the hydrogen bond between N88 Nδ2-Hδ2 and M91 O breaks (5.5 Å), allowing the sidechain of N88 to orient away from the backbone of both K91 and T92 and enabling a more open state (Monomer B: 7.6 Å).

These observations suggest that the sidechain interactions of N88 with the backbones of residues 91 and 92 modulate handle region dynamics: the handle region conformations transition from the most closed structure (Monomer C: 6.7 Å) to the most open structure (Monomer B: 7.6 Å) and vice versa as the N88 sidechain hydrogen bonds break and reform. This mechanism is consistent with previous results showing that the interaction between R88 and T92 was characterized by the handle distance metric with shorter distances more prevalent in closed conformations.^12^ A major difference between the RNHI homologs with Arg or Asn at position 88, however, is that the R88 sidechain is only involved in one interaction, probably water-mediated, with T92 O, whereas the N88 sidechain is involved in up to three hydrogen bond interactions with the handle loop backbone (N88 Oδ1: K91 N-H, N88 Nδ2-Hδ2: K91 O, and N88 Nδ2-Hδ2: T92 O). If all three interactions modulate handle distance, then eight (2^3^) possible handle region conformations can be characterized by the different hydrogen bonding patterns.

Table 1 shows the average handle distances observed when hydrogen bonds are present or absent in MD simulations of the N88 RNHI variants. The handle distances do not depend significantly on the presence or absence of N88 sidechain interactions with the K91 backbone. However, shorter average handle distances are observed when the hydrogen bond between N88 Nδ2-Hδ2 and T92 O is present than when the hydrogen bond is absent (paired t-test, *p* = 0.0097), consistent with data presented so far. Trends are also seen across the homologs for the eight possible combinations of the N88 sidechain interactions (Table 1). Ordering of the eight conformations can be done in similar manner as for the monomers of 6X SoRNHI, as sequential increases in handle distance result in either the loss or gain of a hydrogen bond. Interestingly, this ordering results in the first four combinations all containing a hydrogen bond between N88 Nδ2-Hδ2 and T92 O (preferably closed), while the last four do not (preferably open). As such, the combinations can be grouped into proposed closed conformers that contain this interaction (Figure 6) and open conformers that do not (Figure 7).

**Table 1.** Handle Distance averages of N88 Hydrogen Bonding interactions.

| Protein | 88 Oδ1 : 91 N |  | 88 Nδ2 : 91 O |  | 88 Nδ2 : 92 O |  |
| --- | --- | --- | --- | --- | --- | --- |
|  | Present | Absent | Present | Absent | Present | Absent |
| **R88N EcRNHI** | 8.4(43.9%) | 8.3(56.1%) | 8.3(67.9%) | 8.4(32.1%) | 7.5(12.2%) | 8.4(87.8%) |
| **K88N SoRNHI** | 8.4(57.1%) | 8.5(42.9%) | 8.4(78.2%) | 8.7(21.8%) | 7.7(16.8%) | 8.6(83.2%) |
| **CtRNHI** | 7.6(95.3%) | 7.2(4.7%) | 7.6(97.1%) | 7.9(2.9%) | 6.9(2.9%) | 7.3(93.4%) |
| **HsRNHI** | 8.0(93.2%) | 7.1(6.8%) | 7.9(95.4%) | 7.9(4.6%) | 7.0(7.1%) | 8.0(92.9%) |

| Protein | 88 Oδ1 & 92 O | 92 O | 91 O & 92 O | All | 88 Oδ1 & 91 O | 91 O | None | 88 Oδ1 |
| --- | --- | --- | --- | --- | --- | --- | --- | --- |
| **R88N EcRNHI** | (0%) | 6.5(0.5%) | 7.5(10.2%) | 7.9(1.6%) | 8.4(41.4%) | 8.5(14.8%) | 8.5(30.6%) | 9.1(1.0%) |
| **K88N SoRNHI** | 5.9(<0.1%) | 6.5(0.8%) | 7.7(11.5%) | 7.9(4.5%) | 8.4(50.6%) | 8.9(11.6%) | 8.7(19.1%) | 9.2(1.9%) |
| **CtRNHI** | (0%) | 6.1(<0.1%) | 6.3(1.4%) | 7.6(1.4%) | 7.6(93.4%) | 7.3(0.8%) | 7.8(2.3%) | 8.6(0.5%) |
| **HsRNHI** | (0%) | 6.0(0.9%) | 6.8(3.6%) | 7.6(2.6%) | 8.0(88.2%) | 7.8(1.0%) | 8.2(1.2%) | 8.6(2.4%) |

Top: 100 ns MD simulations at 300 K for 4 RNHI homologs and mutants (R88N EcRNHI, K88N SoRNHI, CtRNHI, and HsRNHI) were separated respectively by the presence or absence of the three hydrogen bonding interactions of the N88 sidechain (N88 Oδ1: K91 N-H, N88 Nδ2-Hδ2: K91 O, and N88 Nδ2-Hδ2: T92 O). The VMD H-Bond plugin (4.0 Å and 120° cutoff) was used to assess the presence of a hydrogen bond. The handle distance averages (Å) of each group and their respective percentages in parenthesis are reported. Bottom: Same as top but frames were separated by the eight possible combinations of interactions. The acceptor atom of each interaction is shown in the column name.

**Figure 6.**
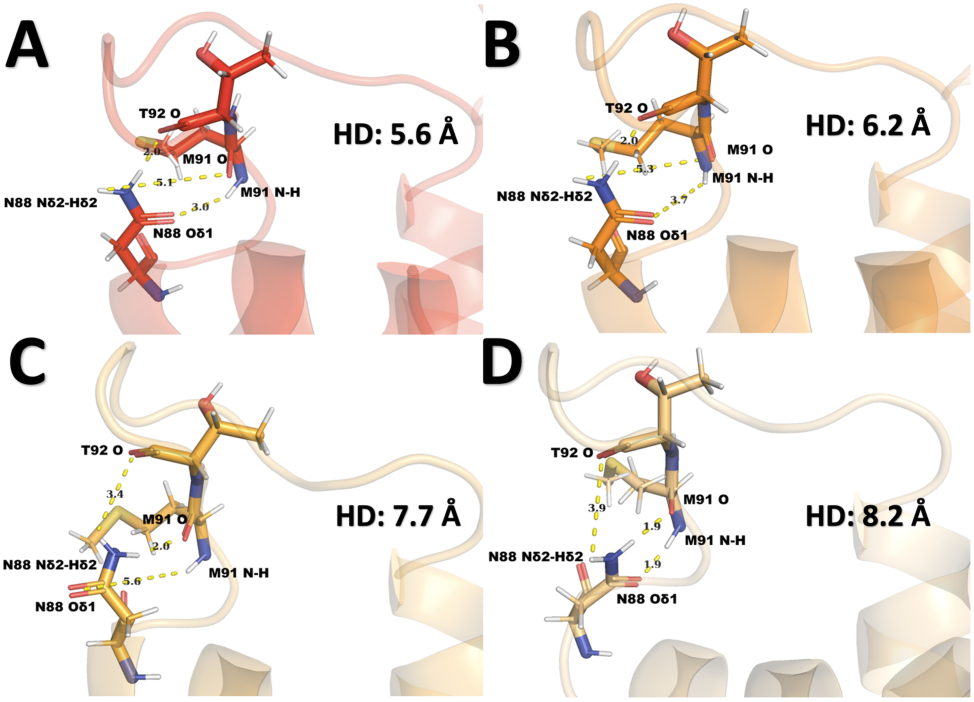
Proposed Closed Conformers for RNHI Homologs. A randomly selected frame was taken from each of the first 4 interaction groups of Table 1B from K88N SoRNHI; K88N SoRNHI is the only RNHI variant that populates all eight interaction combinations in MD simulations. Distances of all 3 potential N88 sidechains are measured and handle distances are also displayed for interaction combinations: **(A)** N88 Nδ2-Hδ2: T92: O and N88 Oδ1: M91 N-H **(B)** N88 Nδ2-Hδ2: T92 O **(C)** N88 Nδ2-Hδ2: T92: O and N88 Nδ2-Hδ2: M91 O **(D)** N88 Nδ2-Hδ2: T92: O, N88 Nδ2-Hδ2: M91 O, and N88 Oδ1: M91 N-H. This group is defined by the presence of N88 Nδ2-Hδ2: T92 O hydrogen bond.

**Figure 7.**
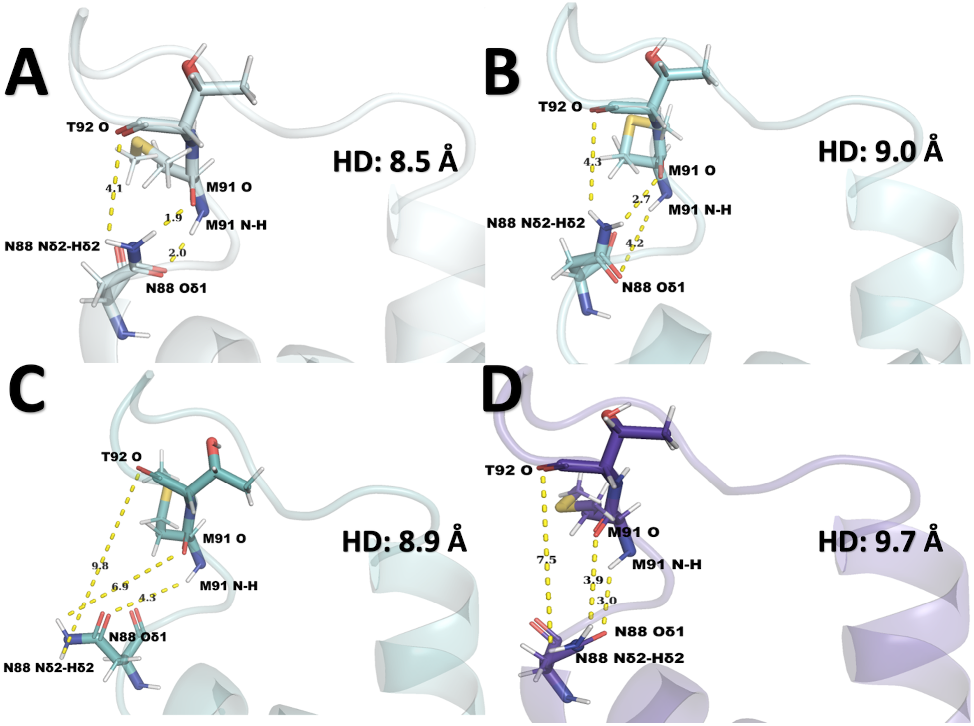
Proposed Open Conformers for RNHI Homologs. Same as Figure 6 but for interaction groups: **(A)** N88 Nδ2-Hδ2: M91 O and N88 Oδ1: M91 N-H **(B)** N88 Nδ2-Hδ2: M91 O **(C)** No sidechain interactions **(D)** N88 Oδ1: M91 N-H. This group is defined by the absence of N88 Nδ2-Hδ2: T92 O hydrogen bond.

## DISCUSSION

CtRNHI has structural and thermodynamic features that are similar to its mesophilic (EcRNHI) and thermophilic (TtRNHI) counterparts. Additionally, MD simulations found that CtRNHI has a one-state handle region conformation that is intermediate between the two-state open and closed handle region conformations of EcRNHI and TtRNHI. To investigate these differences, NMR chemical shifts, scalar coupling constants and RDCs were measured to characterize the conformational properties of the CtRNHI handle region. The weighted conformer score for CtRNHI is intermediate between EcRNHI and TtRNHI, giving a *de novo* K_M_ prediction that is ~2.4 fold greater than EcRNHI; in contrast, activity assays yielded a K_M_ for CtRNHI that is statistically indistinguishable from EcRNHI. This discrepancy is consistent with MD simulations suggesting that CtRNHI populates a state of the handle region in between the open and closed states. RNHI homologs in which position 88 is an Arg residue populate both open and closed states of the handle region in solution and substrate affinity is correlated with the open state population. In contrast, RNHI homologs with Asn at position 88 populate one-state handle region conformations preferentially and behave differently than homologs with two-state handle region conformations. Thus one- and two-state RNHI homologs require different models to describe conformational dynamics and substrate affinity: for CtRNHI the open-state population is not a dominant predictor of substrate affinity.

Comparison of reciprocal mutants at sequence position 88 for EcRNHI and CtRNHI provides insight into the structural features of one-state handle regions. The effect of mutation is predominantly local (confined to the handle region) as evident by ^15^N and ^13^C HSQC chemical shift perturbations for N88R CtRNHI, but small and broadly distributed in R88N EcRNHI. Complementary CD spectroscopy shows that the secondary structures of the homologs are barely changed by mutation. N88 is preferred in thermophilic homologs^10^ and is thermostabilizing with a low mutability score in the psychrophile, SoRNHI.^22^ Furthermore, previous studies of chimeras of RNHI homologs^23–24^ have shown that the hydrophobic core (residues 43–122) correlates with capacity for reversible thermal denaturation: the cores of CtRNHI and TtRNHI are sufficient for renaturation, while the core of EcRNH is insufficient without the use of a chemical denaturant^25^ or low pH.^26^ HsRNHI, a homolog with N88, also unfolds reversibly upon thermal denaturation.^27^ Additionally, N88 containing ancestral mutants of EcRNHI and TtRNHI, with the exception of AncB (last mesophilic ancestor with N88), unfolded reversibly upon thermal denaturation.^28^ These data reinforce the notion of N88 as a key contributor to the thermostability of RNHI enzymes.

Further comparison of N88 RNHI homologs/mutants with R88 RNHI homologs/mutants amplify structural contributions to handle region dynamics.^12^ Arginine has a long guanidino sidechain with only one of its protons in range to interact with the backbones of K91 or T92; a hydrogen bond is formed with K91, but its weaker interaction with T92 might be mediated through a potential water molecule. Asparagine on the other hand has a short carboxyamide sidechain in range to form three hydrogen bonds with backbone atoms of K91 and T92; 2 with K91 and one with T92.

An Asn at position 88 in RNHI homologs can make three side chain hydrogen bonds with residues 91 and 92: N88 Oδ1: K91 NH, N88 Nδ2-Hδ2: K91 O, and N88 Nδ2-Hδ2: T92 O. The conformational states of the handle region for N88 RNHI homologs are modulated by the eight possible combinations of the N88 hydrogen bonds. Four closed states are associated with the presence of the N88 Nδ2-Hδ2: T92 O hydrogen bond (Figure 6) and four open states are associated with the absence of this hydrogen bond (Figure 7). Seven of these eight conformational states are seen among the multiple monomers of 6X SoRNHI (2ZQB) and HsRNHI (2QKK). Moreover, MD simulations of N88 homologs and mutants show that structural transitions of the handle region between consecutive frames change are accompanied by changes in hydrogen bond interactions of ± 1, for a total of 12 unique transitions between ordered states. In this model, the (rare in MD simulations) fully closed state is shown in Figure 6A, in which the N88 Nδ2-Hδ2: T92 O and N88 Oδ1: K91 NH hydrogen bonds are formed. Upon losing the hydrogen bond with K91 NH, the enzyme transitions to the closed state that is most similar to that observed in R88 containing homologs (Figure 6B). Formation of subsequent hydrogen bonds with the backbone of K91, first N88 Nδ2-Hδ2: K91 O and then re-forming N88 Oδ1: K91 NH, yield a pair of closed intermediates (Figure 6C & D). The loss of the hydrogen bond with T92 O yields a pair of quasi-open conformations with both N88 Nδ2-Hδ2: K91 O and N88 Oδ1: K91 NH hydrogen bonds (Figure 7A) or only the N88 Nδ2-Hδ2: K91 O hydrogen bond (Figure 7B). All three hydrogen bonds are absent in the open state similar to that of R88 RNHI homologs (Figure 7C). Formation of the N88 Oδ1: K91 NH hydrogen bond yields a fully open state (Figure 7D). The MD simulations of all four N88 RNHI homologs and mutants follow this kinetic model ≥ 99% of the time (Table S4); an overwhelming 81-96% of these transitions feature no change in hydrogen bond interactions, consistent with the stable nature of one-state handle region conformations. An important consequence of this model is that intermediate open conformations of CtRNHI must have altered affinities for substrate compared with the open conformation to account for the difference between predicted and measured Michaelis constants using the Weighted Conformer Model developed for two-state RNHI homologs.

## CONCLUSION

Residue 88 was previously shown to be a major player in RNHI handle region dynamics, as the choice of amino acid at this position affects conformational preferences: Arginine and lysine is present in homologs with two-state handle regions, while asparagine is present in homologs with one-state handle regions.^10^ A weighted conformer model was designed and shown to correlate open-state populations with the Michaelis constants for enzyme activity for homologs with two-state handle regions, but this model was not tested for homologs with one-state handle regions.^12^ To better understand the mechanisms of substrate recognition by RNHI homologs with one-state handle regions, this study conducted near complete NMR backbone and sidechain resonance assignments of a representative homolog, CtRNHI, and showed that the structural features underlying conformational change in one-state handle regions differs from two-state counterparts. Structural comparisons of N88 homologs and mutants showed a direct correlation between the interactions of N88 with K91 and T92 and a handle distance metric defining a range of conformations between fully open and fully closed states. Using MD simulations of K88N SoRNHI and direct evidence from four monomers in the x-ray crystal structure of a mutant of SoRNHI containing six stabilizing mutations, including K88N, a kinetic model was developed that includes 12 unique transitions between eight conformations of the handle region for N88 RNHI homologs. In contrast to RNHI homologs with two-state handle regions, the Michaelis constant of CtRNHI is not strongly determined by measures of the open state population, implying that other intermediate states make substantial contributions to substrate affinity. These findings exemplify the cohesiveness and power in the use of multiple biophysical tools to paint a more comprehensive picture of the linkage between protein dynamics and activity.

## ASSOCIATED CONTENT

### Supporting Information

Experimental procedures, one equation, five figures, and four tables are included in the Supporting Information.

### Accession Codes

EcRNHI (UniProt P0A7Y4), CtRNHI (Uniprot Q93SU7), SoRNHI (UniProt A0A2W5DXL8), and HsRNHI (UniProt O60930). Backbone ^1^H–^15^N backbone assignments are deposited at BioMagResBank: CtRNHI (BMRB 51160) N88R CtRNHI (BMRB 51162) and R88N EcRNHI (BMRB 51163).

## Notes

The authors declare no competing financial interest.

## ACKNOWLEDGMENT

This paper is dedicated to the memory of Arlene Davis. We acknowledge Kate Stafford (Columbia University) for providing MD simulations; Kate Stafford and Tassadite Dahmane (New York Structural Biology Center) for sample preparation; and Jia Ma (Columbia University) for assistance with CD and fluorescence spectroscopy.

