## Supplemental Information for "Comparisons of ribonuclease HI homologs and mutants uncover a multi-state model for substrate recognition"

### I. Methods

#### *Simulations, Revised Handle Distance Metric, & PALES Predicted RDCs*

All MD simulations analyzed in this study were reported previously (*1*). Briefly, simulations were performed using Desmond with the Amber99SB force field and the TIP3P water model. Electrostatics were calculated with the PME method. All simulations used a 2.5 fs inner time step on a 1-1-3 RESPA cycle. The PDB structures 2E4L, 2RN2, 3HO8, 2QK9 were used to initiate trajectories at 300 K. Computational mutagenesis was performed in Maestro version 9.1. Handle-region dynamics were monitored using a reaction coordinate consisting of the Cartesian distance between the a-carbons of residues equivalent to W85 and T92 in EcRNHI. Values of the distance metric ≥ 7.8 Å (distance measurement for wild-type EcRNHI pdb file 2RN2) were considered to reflect an open state, while distances < 7.8 Å were considered to reflect a closed state. Prediction of residual dipolar coupling (RDC) constants, calculation of alignment tensors, and back calculation of RDCs from experimentally derived alignment tensors were performed in PALES according to previous protocols (*2*). All RDCs were normalized by the ratio of the magnitude of the experimentally derived alignment tensor ($M_{AT}$) of EcRNHI to the alignment tensor of the ensemble of interest. For example, a normalized RDC for TtRNHI is given by: $RDC_{TtRNHI}$ * $M_{AT\_EcRNHI}$ / $M_{AT\_TtRNHI}$.

#### *Sample Preparation*

All protein constructs were Cys-free mutants. Protein expression, purification and NMR sample preparation for the CtRNHI, N88R CtRNHI, EcRNHI and R88N EcRNHI were performed as previously described (*3*). NMR samples were [U-$^{13}$C, U-$^{15}$N] protein in a buffer consisting of 50 mM deuterated sodium acetate, 50 mM sodium chloride, 10% (v/v) deuterium oxide, 0.02% (w/v) sodium azide, and 3 mM sodium 2,2-dimethyl-2-silapentane-sulfonate (DSS) at pH 5.5.

#### *CD Spectroscopy*

CD spectroscopy was performed according to literature (4), using a Chirascan V100 Spectrometer (Columbia Precision Biomolecular Characterization Facility, PBCF). Spectra were collected using 1mg/mL protein samples in a buffer solution of 20 mM sodium acetate, 50 mM potassium chloride at pH 5.5 in a 0.1 cm quartz cuvette at 25 °C. The far UV range, 250–200 nm, was used to collect data points at 1-nm intervals. Thermal denaturation melts were performed by heating the sample from 25

°C to 85 °C and subsequent cooling back to 25 °C at intervals of 3 °C. Data was collected at each interval. Additionally, spectra were taken at 25 °C before and after the thermal denaturation to test for reversibility.

#### *NMR Spectroscopy*

NMR spectra were recorded using Bruker Biospin Avance 600 MHz (Columbia University) and Avance 800 MHz (New York Structural Biology Center, NYSBC) NMR spectrometers equipped with TCI z-axis gradient cryoprobes. Spectra were processed with nmrPipe (*5*) and analyzed using NMRFAM-Sparky (*6*). Spectra were referenced to DSS for $^{1}H$ resonances and indirectly for $^{13}C$ and $^{15}N$ resonances (*7*).

#### *Backbone and Methyl Group Assignments*

Backbone and methyl group assignments have been reported for EcRNHI (*8*). Backbone ($^{1}H$, $^{15}N$, $^{13}C_{\alpha}$, $^{13}C'$) and sidechain ($^{13}C_{\beta}$) resonance assignments for CtRNHI, N88R CtRNHI and R88N EcRNHI were obtained using 3D $^{1}H$-$^{15}N$ NOESY-HSQC, TROSY HNCA, HN(CO)CA, HNCACB, HN(CO)CACB, HNCO NMR experiments (*9*). Aliphatic sidechains were assigned using 3D $^{1}H$- $^{13}C$ NOESY-HSQC, HC(C)H-COSY, and (H)CCH-COSY experiments (*10-12*). Stereospecific assignments of valine and leucine sidechains were assigned using samples fractionally labelled with 10% $^{13}C$ (*13-14*).

#### *Residual Dipolar Coupling Constants*

Samples in anisotropic media were prepared as previously described (*15*). Briefly samples were partially aligned by mixing 100 µl of 16% Otting media (50 µl $C_{12}E_5$ (pentaethylene glycol monododecyl ether, PEG), 200 µl NMR sample buffer, 50 µl of $D_2O$, 16 µl hexanol) with 300 µl of the isotropic protein sample to give partial alignment with a final PEG concentration of 4%. Triplicate IPAP experiments (*16*) were recorded for both the aligned and isotropic samples of each protein at 18.8 T and 298 K. Frequency differences between doublet components were measured for the isotropic and aligned samples. The RDCs were obtained from the difference in splittings between the aligned and isotropic samples.

#### *Scalar Coupling Constants and Rotamer Populations*

Three-bond $^{13}C'$-$^{13}C\gamma1$, $^{13}C'$-$^{13}C\gamma2$, $^{15}N$-$^{13}C\gamma1$ and $^{15}N$-$^{13}C\gamma2$ scalar coupling constants were measured at 14.1 T and 300 K (*17-18*). Rotamer distributions were calculated from the scalar couplings using established models (*19*) and from methyl group $^{13}C$ chemical shifts using the SideR program (*20*). Dihedral angles were for individual frames of MD simulations were determined using VMD (*21*) and aggregated to give populations of gauche+, gauche–, and trans rotamers.

#### *Enzyme Kinetic Assays*

Comparative binding assays for EcRNHI and CtRNHI were performed following an established method (*22*). Briefly, reactions were performed in a solution of 10 mM Tris–HCl, 50 mM NaCl, 10 mM $MgCl_2$, pH 8 at 25 °C. The substrate was RNA–DNA hybrid labeled with a fluorophore-quencher pair (RNA = Fl-5'-GAGAUGACGG-3', DNA = 5'-CCGTCATCTC-3'-DA, Fl = 5' 6-FAM (fluorophore: Absorbance Max- 495 nm, Emission max- 520), DA =DABCYL (quencher)). The hybrid concentration was varied from 0 nM to 600 nM. Fluorescence was monitored using a Horiba Lifetime Fluorimeter (Columbia PBCF) for the reaction mixture containing only substrate for 1 minute to establish baseline emission. Fluorescence monitoring was paused to add enzyme to a final concentration of 3 nM EcRNHI or CtRNHI. Fluorescence monitoring was resumed for 10 minutes. Hydrolysis of the RNA strand leads to an increase in fluorophore emission as a function of time after mixing enzyme and substrate owing to increased separation from the quencher.

#### *Weighted Conformer Score*

The Weighted Conformer Score for each protein is calculated from the experimental values of W85 Nε1-Hε1 RDCs, T92 N-H RDCs and residue 101 trans rotamer percentages using:

$$\textbf{Weighted Conformer Score} = \{(W85CS*r^2_{W85}) + (T92CS*r^2_{T92}) + (Trans\%*r^2_{Trans})\} / 3 \qquad (S1)$$

Trans% is the average percentage of trans population for residue 101 calculated from each method, reported from 0 to 1. W85CS and T92CS are the conformer scores assigned to each respective RDC. The highest and lowest RDC values of the predicted RDC distributions (95% of the values in the center of the distribution) were assigned scores of 0 (completely open) and 1 (completely closed) respectively. The experimental RDC value was given a score based off its position in this range. Each parameter in Eq. S1 for a given RNHI protein is weighted by the Pearson's correlation coefficient ($r^2$) determined from a plot of normalized $K_M$ values versus parameter for the full set of RNHI proteins.

**II. Supplemental Figures**

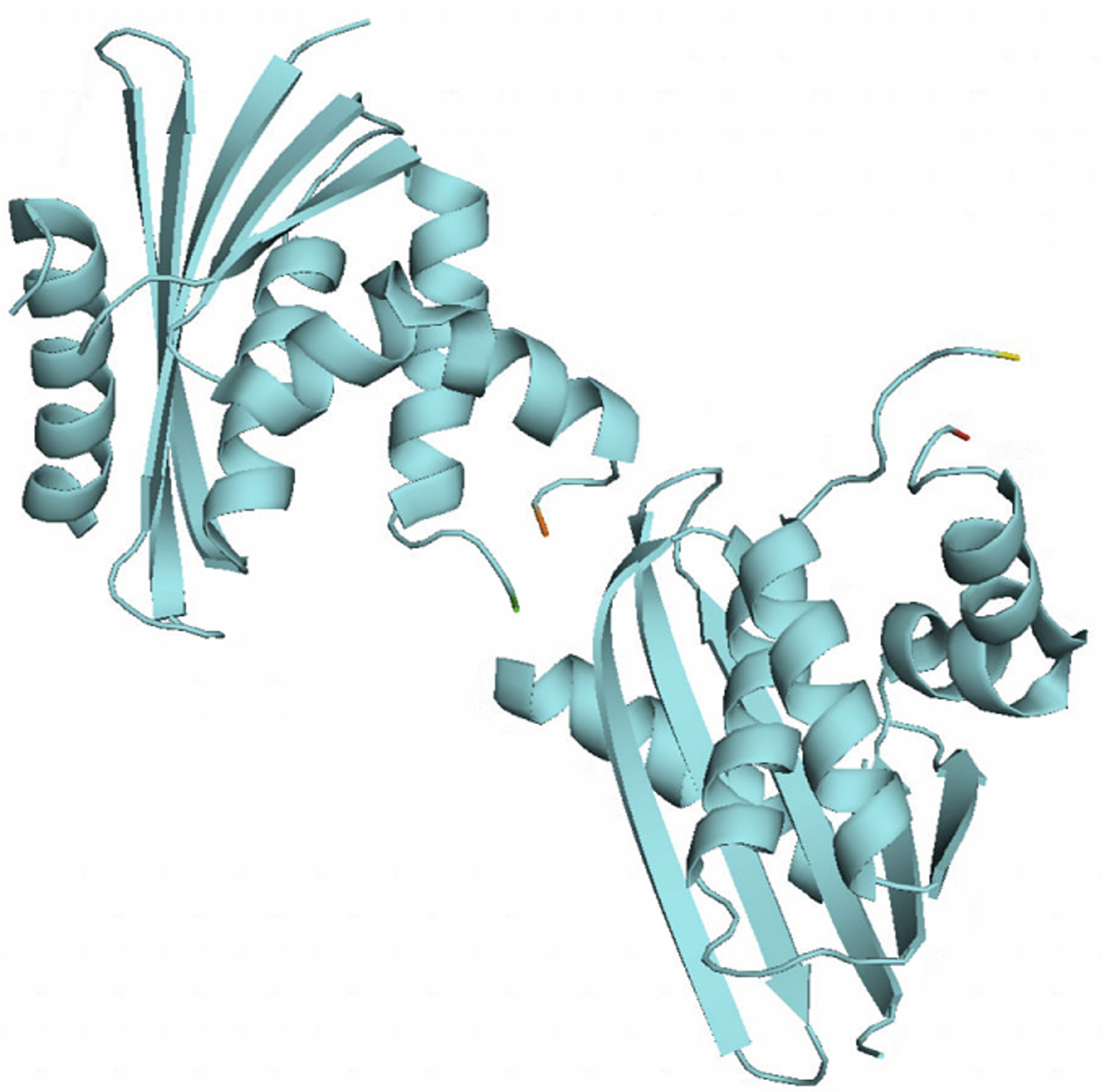

**Figure S1. Tertiary structure of CtRNHI.** X-ray crystallography structure of CtRNHI (PDB ID: 3H08). The unit cell contains two monomers with a Cα-RMSD of 0.44 Å. Each monomer has missing residues in the handle region. The residues preceding the missing ones are color coded as follows: orange (K91), Green (K96), Red (T92), Yellow (A94).

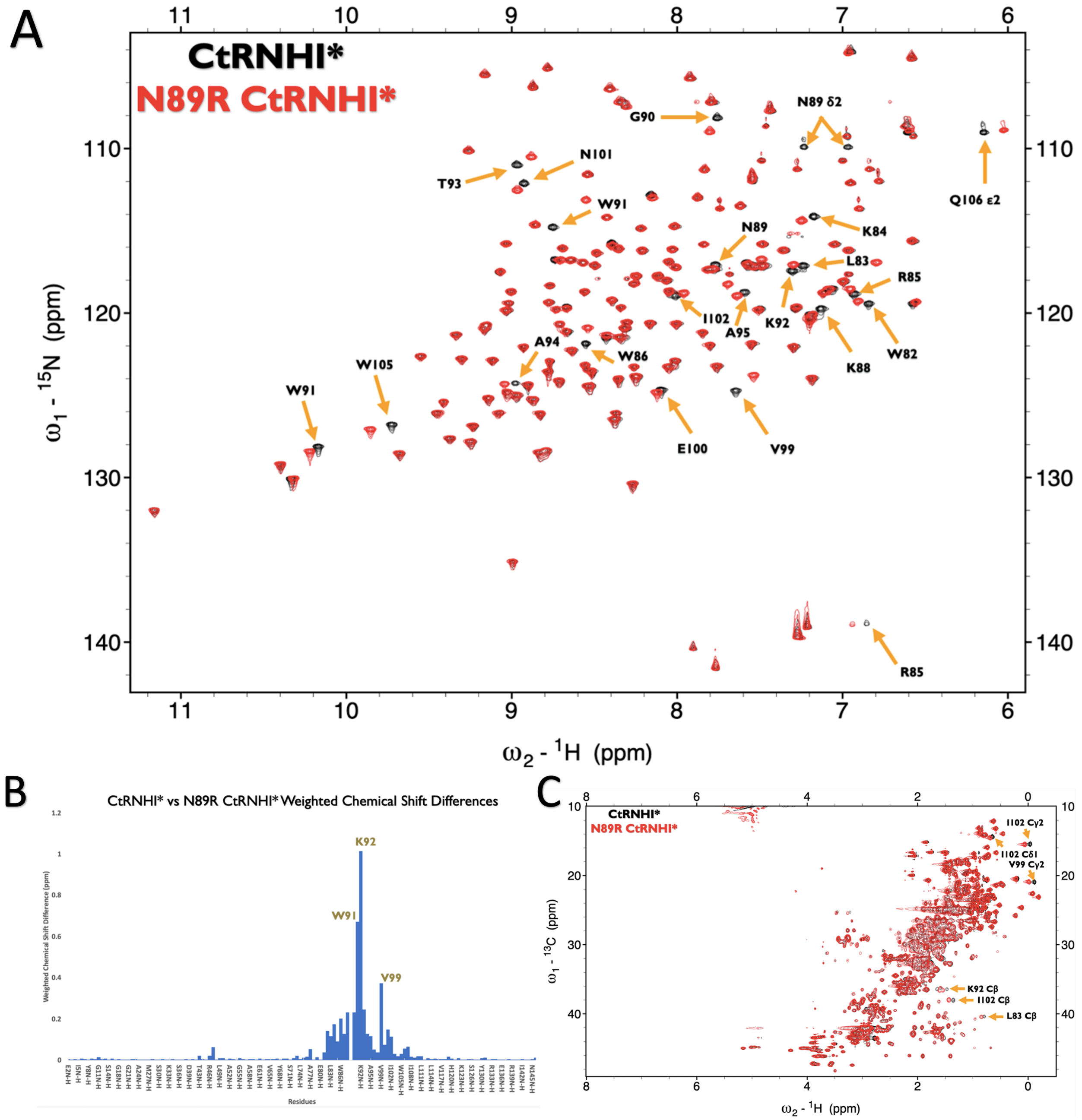


**Figure S2. Wt. vs. N89R CtRNHI Backbone and side chain Comparisons. (A)** Overlaid $^{15}$N HSQC NMR spectra of Wt. (black) & N89R CtRNHI (red). Labeled peaks are for Wt. CtRNHI*. **(B)** Weighted chemical shift differences of the spectra in A. (C) Overlaid $^{13}$C HSQC NMR spectra of Wt. (black) and N89R CtRNHI* (red). Labeled peaks are for Wt. CtRNHI.

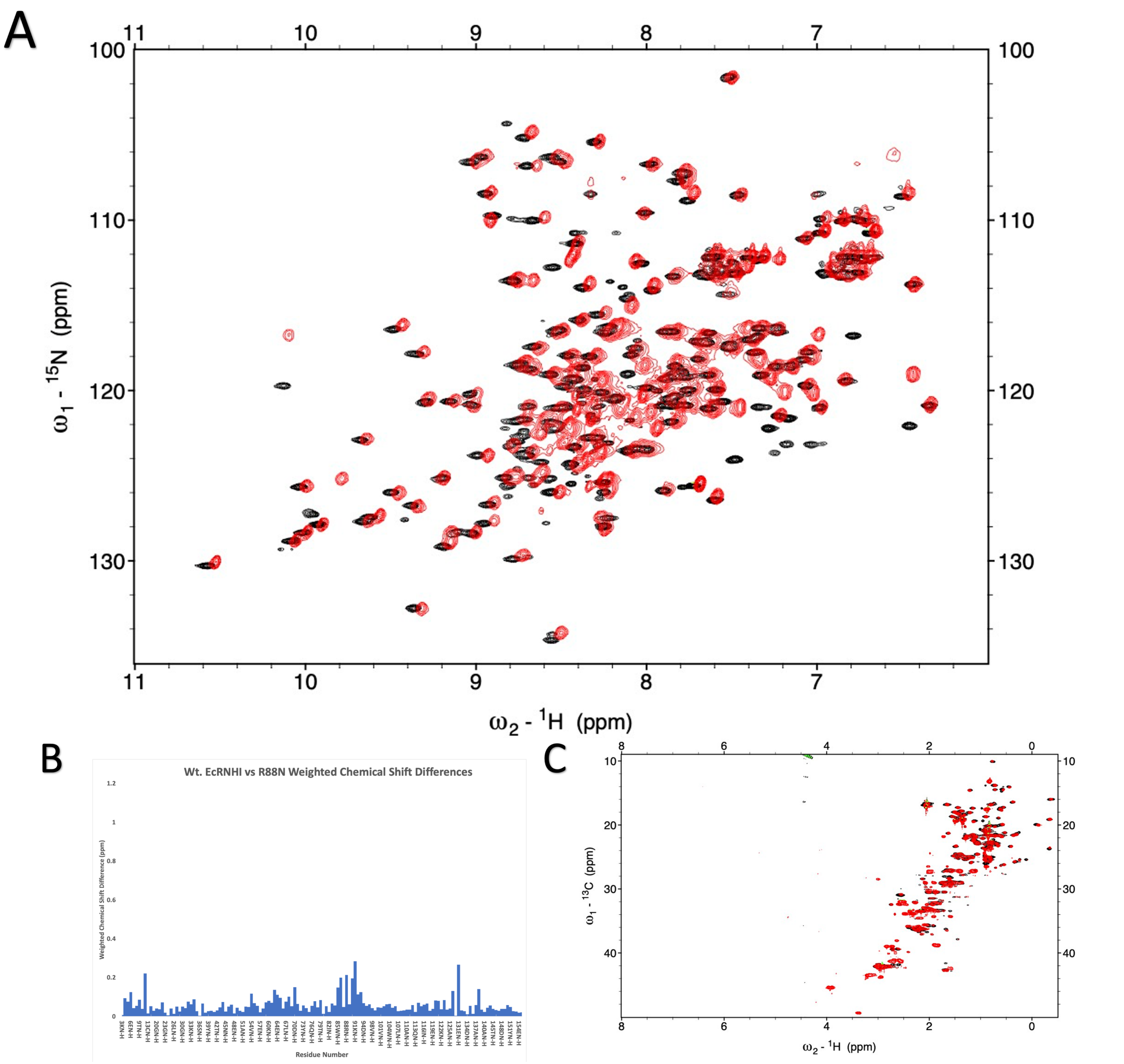


**Figure S3. Wt. vs. R88N EcRNHI Backbone and side chain Comparisons.** Same as Figure S2 but for Wt. EcRNHI (Black) and R88N EcRNHI (Red).

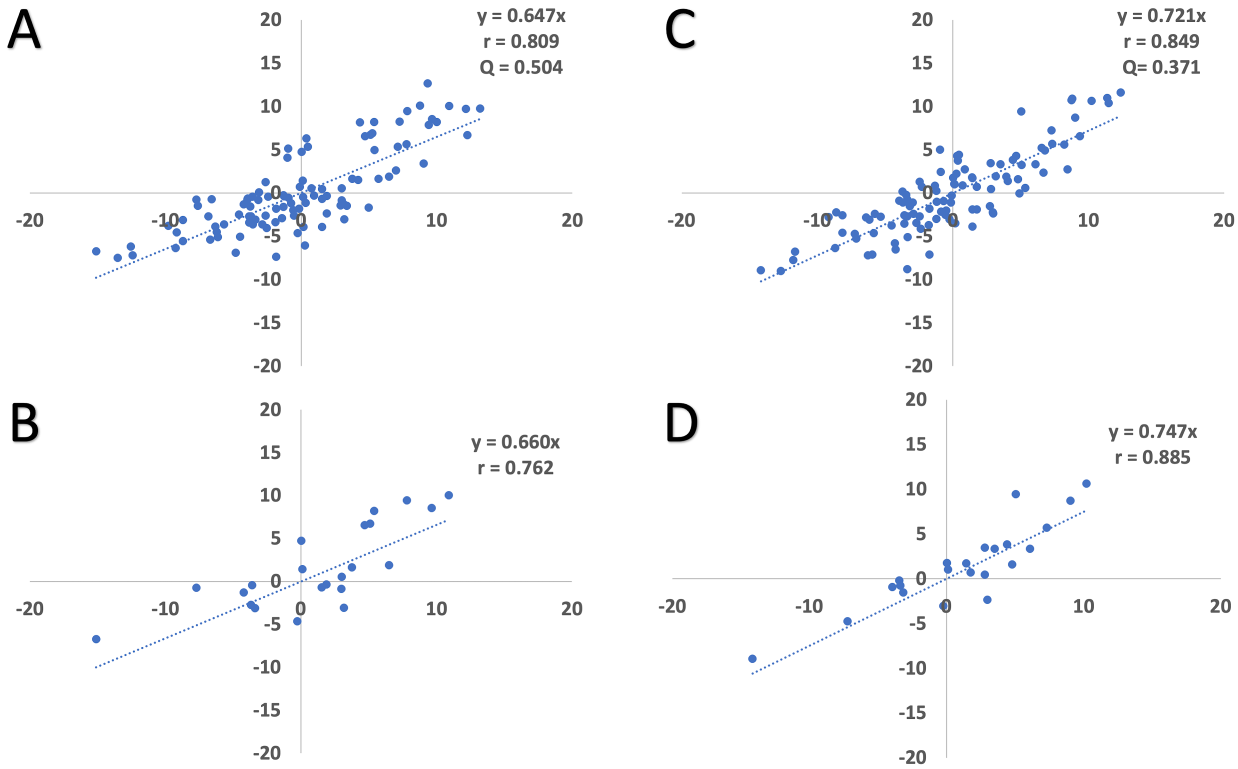


**Figure S4. CtRNHI* Experimental RDCs.** Experimental RDCs (Hz) of CtRNHI* plotted along the x-axis against **(A)** best fitted RDCs of its crystal structure (3H08). **(B)** Same as previous but only the handle region. (**C-D**) Same as A-B but for its best-fit frame in its MD simulations.

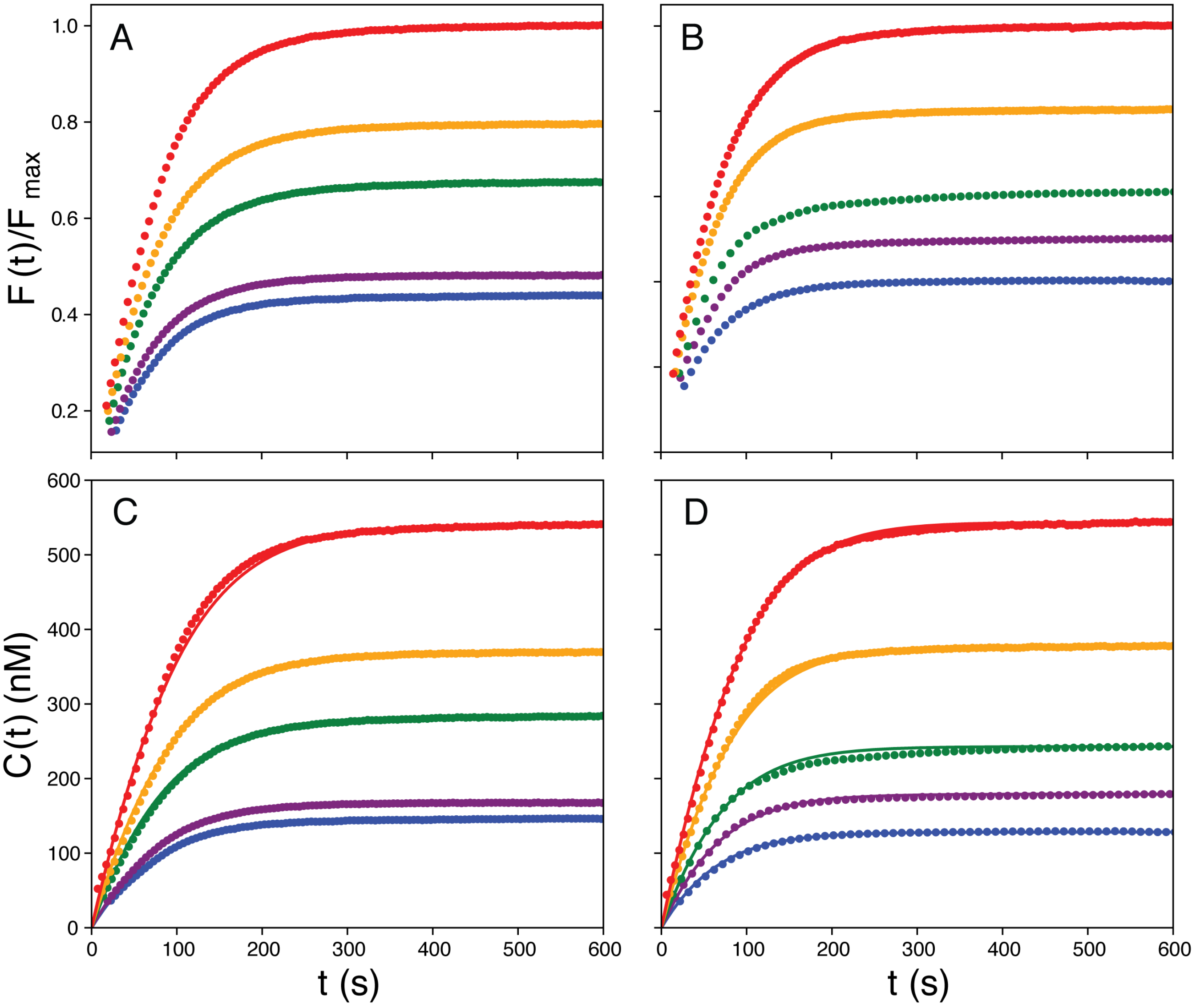


**Figure S5. Enzyme kinetics for CtRNHI and EcRNHI.** Fluorescence time courses for (A) EcRNHI with substrate concentrations of (blue) 146 nM, (purple) 182 nM, (green) 268 nM, (orange) 385 nM, and (red) 518 nM and (B) CtRNHI with substrate concentrations of (blue) 129 nM, (purple) 175 nM, (green) 261 nM, (orange) 397 nM, and (red) 522 nM. Data are normalized the limiting fluorescence after the reaction ran to completion for the largest substrate concentration. Fluorescence values were converted to product concentrations C(t) for reactions with (C) EcRNHI and (D) CtRNHI using limiting values of the fluorescence signal after the reaction ran to completion and depicted as circles. Fluorescence was corrected for quenching effects. The progress curves were globally fit to the integrated Michaelis-Menton equation. Fitted results are shown as solid lines. Data for a substrate concentration of 261 nM for CtRNHI was sampled less frequently than for other conditions and could not be fit stably. This progress curve was excluded from the global fit, but the calculated curve is shown for illustration. Uncertainties in $K_M$ and $v_{max}$ were obtained by a jackknife simulation in which one progress curve at a time was omitted from analysis.

### III. Supplemental Tables

#### Table S1. Enzyme kinetic data and Weighted conformer scores of RNHI homologs.

| Protein | T92 RDC (Hz) | W85 RDC (Hz) | Trans % | WCS | $K_M$ ratio* |
|---|---|---|---|---|---|
| Wt. EcRNHI | -1.47 (0.66) | 3.61 (5.07) | 0.11 | 0.21 | 1.00 |
| V98A EcRNHI | -3.57 (-3.77) | -0.87 (-3.01) | 0.43 | 0.43 | 3.73 |
| SoRNHI | (-3.09) | -0.67 (-7.87) | 0.16 | 0.33 | 2.73 |
| TtRNHI | -8.21 (-6.27) | (-4.86) | 0.85 | 0.63 | 7.36 |
| CtRNHI | -3.70 (-2.51) | -0.02 (5.09) | 0.24 | 0.31 | 0.91 |

Weighted conformer scores for each homolog and mutant were calculated with equation S1 using experimental RDC values for T92 N-H, W85 Nε1-Hε1 and average trans % of residue 101; RDCs back calculated from the respective experimentally derived alignment tensors are in parenthesis. Based off CtRNHI's weighted conformer score, an estimate of its $K_M$ ratio (prediction indicated by asterisk) to Wt. EcRNHI was back calculated from the equation of fit of the other homologs. *$K_M$ ratios are taken from a previous study[23] and calculated according to the following: Homolog $K_M$/Wt. EcRNHI $K_M$, except for CtRNHI measured in the present study.

#### Table S2. Geometric analysis of the crystal and *in silico* structure of CtRNHI

| Geometry Evaluation | 3H08:A | 3H08:B | Bf CtRNHI |
|---|---|---|---|
| # missing residues | 7 | 4 | 0 |
| Clashscore (percentile) | 4.36(95$^{th}$) | 10.55(68$^{th}$) | 0(100$^{th}$) |
| Poor rotamers (%) | 4.17 | 1.69 | 1.83 |
| Favored rotamers (%) | 94.17 | 95.76 | 95.41 |
| Ramachandran outliers (%) | 0 | 0 | 0.74 |
| Ramachandran favored (%) | 100 | 100 | 96.3 |
| MolProbity score (percentile) | 1.61(92$^{nd}$) | 1.72(89$^{th}$) | 0.95(100$^{th}$) |
| Bad bonds (%) | 0 | 0.09 | 0.18 |
| Bad angles (%) | 0 | 0 | 0.85 |

The two monomers in the crystal structure (3H08) and best fitted *in silico* structure of CtRNHI were analyzed using Molprobity and the evaluation criteria are reported. Clashscore represents the number of serious steric overlaps (> 0.4 Å) per 1000 atoms and Molprobity score is a combination of the clashscore, rotamer and Ramachandran evaluations. Smaller Molprobity scores reflect better geometry.

**Table S3. Comparison of Handle region metrics between Wt. & 6X SoRNHI.**

| Protein | PDB | Handle Distance (Å) | V98 χ1 (°) | V101 χ1 (°) | All Ca-RMSD (Å) | Handle region Ca-RMSD (Å) |
|---|---|---|---|---|---|---|
| Wt. SoRNHI | 2E4L | 7.5 | 161.2 | 55.4 | - | - |
| 6X SoRNHI | 2ZQB:A | 6.8 | 175 | 74.2 | 0.653 | 0.357 |
| | 2ZQB:B | 7.6 | -43.4 | 53.9 | 0.926 | 0.904 |
| | 2ZQB:C | 6.7 | 173.9 | 70.0 | 0.906 | 0.608 |
| | 2ZQB:D | 7.4 | 172.6 | 70.3 | 1.062 | 0.817 |

Handle region metrics and structural comparison of Wt. SoRNHI with each monomer of 6X SoRNHI. Both the overall structure and the handle region of each monomer was aligned to Wt. SoRNHI and the Cα-RMSDs are reported.

**Table S4. Change in N88 sidechain hydrogen bond interactions along MD trajectories.**

| Protein | ΔH-bond = 0 | ΔH-bond = 1 | ΔH-bond = 2 | ΔH-bond = 3 |
|---|---|---|---|---|
| R88N EcRNHI | 85.07% | 13.94% | 0.99% | 0.00% |
| K88N SoRNHI | 81.09% | 17.82% | 1.08% | 0.01% |
| CtRNHI | 95.68% | 4.19% | 0.13% | 0.00% |
| HsRNHI | 89.86% | 9.70% | 0.44% | 0.00% |

The change in the hydrogen bond interaction between consecutive frames of the three sidechain atoms of N88 and the backbones of the handle loop (K91 & T92) were mapped along 100 ns trajectories at 300K for 4 RNHI variants (R88N EcRNHI, K88N SoRNHI, CtRNHI and HsRNHI). The absolute sum of the three interaction changes is shown with ΔH-bond=0 indicating that the interactions at all 3 positions did not change between frames, ΔH-bond=1 indicating that an interaction at one of the three positions was either gained or lost between frames, so on so forth. Percentages of each group are reported for each RNHI variant. The VMD H-Bond plugin (4.0 Å and 120° cutoff) was used to assess the presence of a hydrogen bond.
